# Predicting *Blomia tropicalis* allergens using a multiomics approach

**DOI:** 10.1101/2023.05.10.540119

**Authors:** Jan Hubert, Susanne Vrtala, Bruno Sopko, Scot E. Dowd, Qixin He, Pavel B. Klimov, Karel Harant, Pavel Talacko, Tomas Erban

## Abstract

The domestic mite *Blomia tropicalis* is a major source of allergens in tropical and subtropical regions. Despite its great medical importance, the allergome of this mite is not sufficiently studied. Only 14 allergen groups have been identified in *B. tropicalis* so far, even though early radioimmunoelectrophoresis techniques (27 uncharacterized allergen complexes) and comparative data based on 40 allergen groups officially recognized by WHO/IUIS in domestic astigmatid mites, suggest that a large set of additional allergens may be present. Here, we use a multiomics approach to assess the allergome of *B. tropicalis* using genomic, transcriptomic sequence data, and a highly sensitive protein abundance quantification. Out of 14 known allergen groups, we confirmed 13 (one WHO/IUIS allergen, Blo t 19, was found to be a contaminant originated from a nematode) and identified 16 potentially novel allergens using sequence similarity. These data indicate that *B. tropicalis* shares 27 known/deduced allergen groups with pyroglyphid house dust mites (genus *Dermatophagoides*). Among them, five allergen-encoding genes were highly expressed at the transcript level: Blo t 1, Blo t 5, Blo t 21 (known), Blo t 15, and Blo t 18 (predicted). However, at the protein level, a different set of most abundant allergens was found: Blo t 2, 10, 11, 20 and 21 (mite bodies) or Blo t 3, 4, 6 and predicted Blo t 13, 14 and 36 (mite feces). We show the use of an integrated omics method to identify and predict an array of mite allergens and advanced, label-free proteomics to determine allergen protein abundance. Our research identifies a large set of novel putative allergens and shows that expression levels of allergen-encoding genes may not be strictly correlated with the actual allergenic protein abundance in the mite bodies.

## Introduction

House dust mites and domestic mites cause IgE-mediated type I allergic conditions in an estimated 30% of the world population [1]. In sensitized individuals, allergen exposure induces cross-linking of high-affinity FcεRI receptor-bound IgE on effector cells leading to the release of mediators that cause allergic symptoms [2]. In severe cases, mite allergies manifest in chronic diseases, such as allergic rhinitis, conjunctivitis, and bronchial asthma [3]. Currently, 40 allergens of astigmatid mites are recognized by the Allergen Nomenclature Subcommittee of the World Health Organization (WHO) and the International Union of Immunological Societies (IUIS),[4] however, not all domestic astigmatid mite species are evenly studied for their allergens.

*Blomia tropicalis* is one of the most common and important house dust mite species inhabiting human houses in tropical or subtropical areas [5–9]. Within its typical range, this mite species can be found in up to 96% of dust samples with 40% prevalence of all mite species [7]. Two *B. tropicalis* allergens are considered major allergens rBlo t 5 and rBlo t 21 show IgE reactivity to 93 and 89% of sera from adult asthma patients, respectively [10]. Overall, 27 antigen-antibody precipitin complexes have been identified in of *B. tropicalis* by a crossed radioimmunoelectrophoresis [11]. However, out of 40 allergen groups known in domestic mites, only 14 allergens groups have been described in *B. tropicalis* so far [4, 12]. Next 7 is predicted based on their high similarity to identified allergens of other astigmatid mites [12, 13]. These data collectively indicate that *B. tropicalis* is likely to have a substantial number of unrecognized allergen groups.

One of the major challenges in molecular allergy research is accurate prediction of the allergenic potential of proteins [14]. Accurate diagnosis and targeted therapies for patients are critical for identifying allergenic mite species [13]. The diagnosis is focused on single allergens of mites to determine the exact IgE sensitization pattern of a patient and to distinguish between cosensitization and cross-sensitization [13]. Additional challenges associated with research linked to mite allergenicity is the lack of screening tools for quantifying all allergenic proteins in mite bodies and feces. Mite fecal pellets is arguably the most important allergy-inducing component as they contain powerful digestive proteins [15]. The feces to accumulate overtime in human-related environments and, thanks to their minute sizes, can easily get airborne, thus increasing the risk of contact with the skin and lung epithelium when inhaled [15]. Even though only a small portion of the entire mite allergome may be present in mite feces [16], quantification of these allergens is important. Mite feces may have distinct properties affecting sensitization and allergy-inducement in different ways in comparison to mite themselves. Using a proteomic approach, allergen abundance and diversity have been accurately estimated in several mite species, such as *Dermatophagoides farinae*, *Dermatophagoides pteronyssinus*, and *Tyrophagus putrescentiae* [17–19]; however, data on *B. tropicalis* are lacking. Earlier studies employing bottom-up gel-based proteomics relied on narrow subsets of allergen sequences available at those times [20].

In this study, we generated both a high quality *de novo* whole genome (long and short reads) and transcriptome of *B. tropicalis* and annotated them. Putative allergens were identified based on high sequence identities and structural similarities with known allergens [4] and allergen-related proteins available in public databases. We also estimated the expression levels of known allergens/putative allergens in mite bodies and feces. An advanced proteome analysis using an Orbitrap tribrid mass spectrometer, and deduced allergen sequences were then run to quantity allergic protein richness and abundance in mite bodies and feces.

## Results

### Genome, transcriptome and proteome of *B. tropicalis*

*Blomia tropicalis* predicted genome had 1,601 contigs of 61 Mb length (N_50_=223 kB), according to BUSCO prediction to Arachnida set the completeness was estimated to 93.2% with 2.6% of duplicated genes (Genbank Submission SUB7787303). The annotated transcriptome had 7,171 contigs with a total length of 31 Mb (N_50_=5,996 bp), and 18,164 predicted genes. Its completeness was estimated at 95%, with 9.1% duplicated genes based on BUSCO with the Arachnida dataset. In contrast, a previously generated and annotated transcriptome had a fewer predicted genes 16,590-14,899 [21, 22]. The proteome analyses identified 3,714 protein hits in the four protein sample types: individually collected adult mites; pools of mites of different ages, including eggs; water extracts of feces; and detergent-buffer extracts of the remaining pellet of the water extract.

### Identification of known allergens and putative allergens in the mite transcriptome and proteome

We inferred a total of 31 allergen/putative allergen-encoding proteins (Table 1, S1–S50 Figs, S1–S30 Tables). Of them, 13 proteins were confirmed WHO/IUIS allergens (groups 1-8,10-13, 21); 17 proteins represented 16 allergen groups known in other mites (14–16, 18, 20, 24–26, 28–30, 32–35, 36) but not medically evaluated in *B. tropicalis*, and 1 protein (60S acidic ribosomal protein P2) represented by a putative new allergen group [20] (Table 1: nd). Group 36 allergen wrongly included two different structural groups (Table 1: group 36a,b). The groups of allergens (1,2,4,5,7,14,21) showed significant differences in protein identify based on mite species comparison (Table 2). Identity of *B. tropicalis* allergens or putative allergens significantly differed from the rest of mites species in groups 1,2,3,4,5,14,28.

**Table 1.**
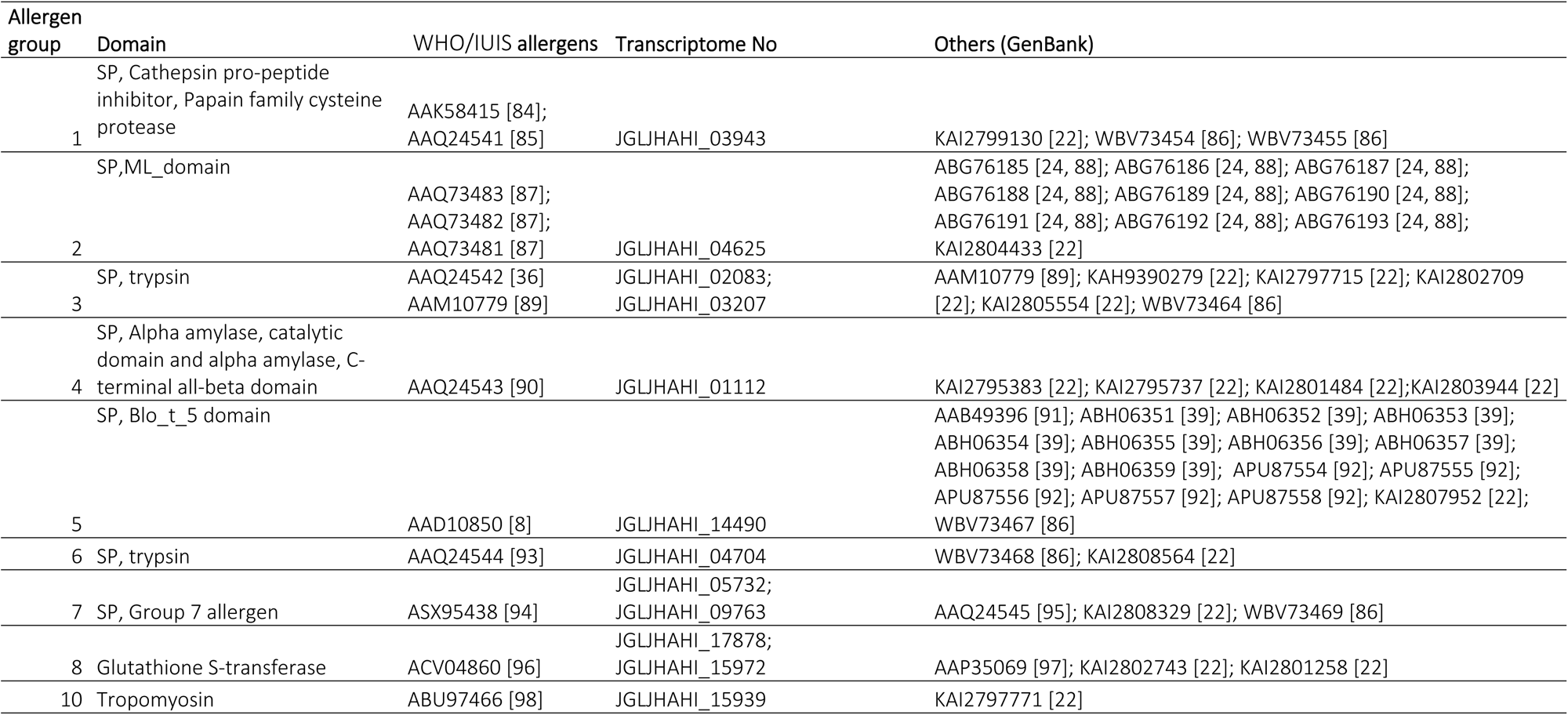
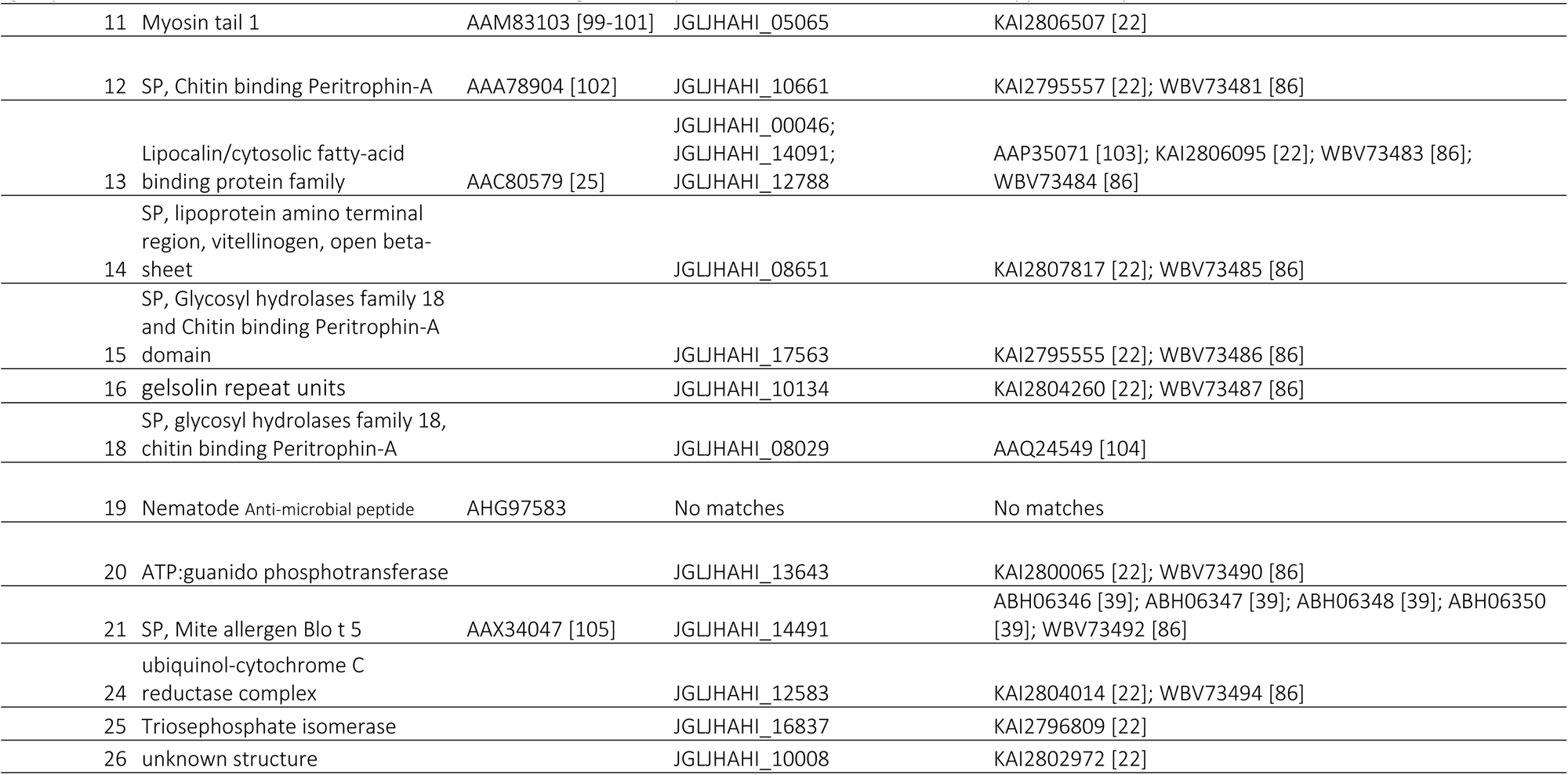
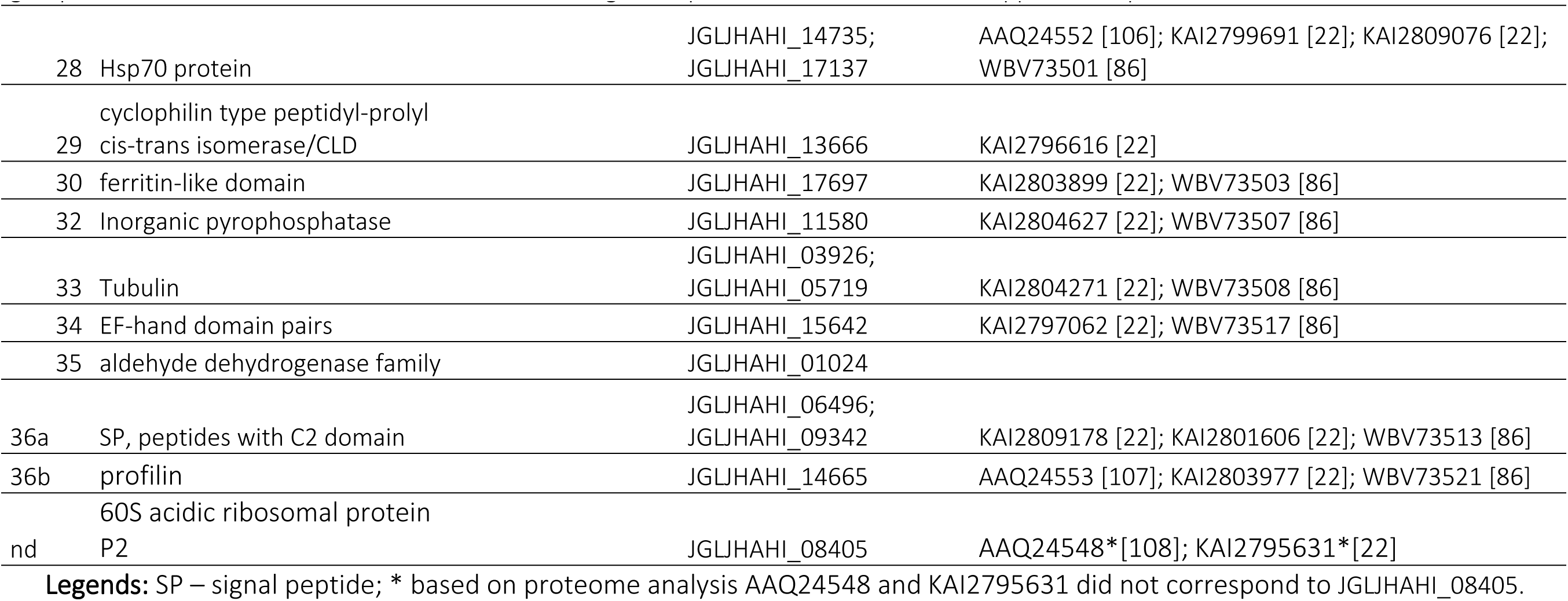
Known and predicted allergen groups in *Blomia tropicalis* based on structural identity analyses (PHMMER) of transcriptomic data. Confirmed allergens from the WHO/IUIS allergen database and closest GenBank matches (annotated or unannotated) are given. Full analysis (allergen and non-allergen proteins) and alignments are given in Supplementary results and Supplementary figures: S1–S50 respectively.

**Table 2.** The comparison of sequences identity based on PERMANOVA (Bray–Curtis, 1000 permutations), the factor were species of mites or *Blomia tropicalis* versus other species. The minimum of sequnces identity is provided and indicated by color scale.

| Factor group | species |  |  | *B. tropicalis* x rest species |  |  |
| --- | --- | --- | --- | --- | --- | --- |
|  | F | P | MIN<sup>#1</sup> | F | P | MIN<sup>#2</sup> |
| 1 | 22.44 | <0.001 | 30.4 | 33.66 | <0.001 | 34.9 |
| 2 | 1291 | <0.001 | 38.5 | 197 | <0.001 | 86.6 |
| 3 | 1.204 | 0.100 | 45.000 | 3.58 | 0.004 | 49.6 |
| 4 | 84.22 | 0.005 | 62.9 | 49.85 | 0.001 | 92.7 |
| 5 | 132.2 | 0.008 | 42.4 | 243.1 | 0.009 | 78.1 |
| 6 | 0.00029 | 0.198 | 55.6 | 8.509 | 0.065 | 99.6 |
| 7 | 5.836 | 0.009 | 24.7 | 8.116 | 0.01 | 43.3 |
| 8 | 1.573 | 0.127 | 32.8 | 4.052 | 0.011 | 34.4 |
| 10 | 0.7165 | 0.689 | 93.3 | 1.006 | 0.4748 | 94 |
| 11 | nd | nd | 86.2 | 3.568 | 0.066 | 91.5 |
| 12 | nd | nd | nd | nd | nd | 91.2 |
| 13 | 0.674 | 0.992 | 51.9 | 0.8726 | 0.409 | 52.1 |
| 14 | 862.1 | 0.017 | 43 | 584.2 | 0.018 | 99.1 |
| 15 | 1696 | 0.052 | 61.4 | 250 | 0.1024 | 98.3 |
| 16 | nd | nd | 56.5 | nd | nd | 99.6 |
| 18 | nd | nd | 60.5 | 39.07 | 0.3235 | 99.6 |
| 20 | 0.4785 | 0.74 | 67.3 | 0.4892 | 0.5234 | 68.3 |
| 21 | 0.00029 | 0.022 | 38.5 | 1180 | 0.0216 | 98.4 |
| 24 | 1794 | 0.1 | 72 | 1794 | 0.1 | 99.2 |
| 25 | 36.64 | 0.203 | 35 | 64.99 | 0.105 | 35 |
| 26 | 0.0008 | 0.161 | 82.6 | 44.3 | 0.333 | 100 |
| 28 | 1.845 | 0.203 | 69.5 | 2.748 | 0.022 | 82.1 |
| 29 | 0.75 | 0.47 | 65 | 0.819 | 0.338 | 65 |
| 30 | 11.68 | 0.099 | 76.5 | 5.409 | 0.099 | 90.8 |
| 32 | nd | nd | 67.4 | 7.871 | 0.103 | 100 |
| 33 | 23.66 | 0.168 | 80.9 | 23.66 | 0.168 | 93.3 |
| 34 | 87.23 | 0.395 | 26.7<sup>#3</sup> | 1.35 | 0.407 | 99.3 |
| 35 | nd | nd | 73.3 | nd | nd | nd |
| 36a | 2.3 | 0.203 | 43.6 | 2.3 | 0.203 | 52.3 |
| 36b | nd | nd | 94.7 | 1.8 | 0.397 | 100 |

**Legend:** nd -not determined, #1 Identity minimum from all sequences; #2 Identity minimum for *B. tropicalis* sequences; #3 The minimum caused by wrongly included BAV90601.

For each of these allergen/putative allergens, protein alignment and phylogenetic trees (*B. tropicalis* and other mites) are given in Supplementary results (S1–S30 Figs). *Blomia tropicalis* had the greatest number of WHO/IUIS allergens shared with pyroglyphid house dust mites *D. farinae* and *D. pteronyssinus* (34%) as opposed to the mold mite *T. putrescentiae*. Twenty-six allergens of *B. tropicalis* were shared with those of pyroglyphid house dust mite genus *Dermatophagoides* mites (Fig 1).

**Fig 1.**
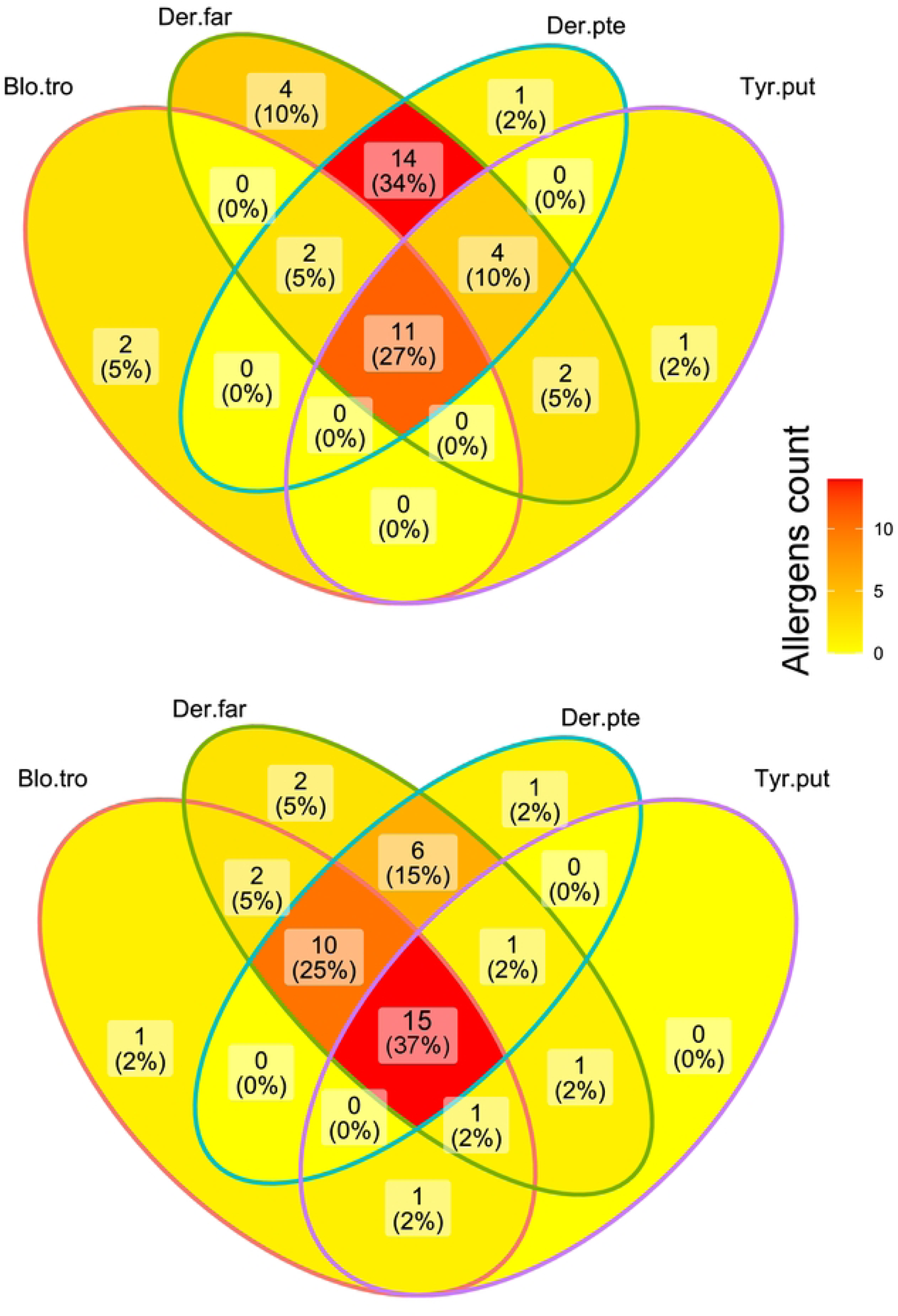
Venn diagrams of allergens from the WHO/IUIS database only (**A**) and WHO/IUIS allergens plus predicted allergens (**B**) in four species of mites: *Dermatophagoides farinae*, *Dermatophagoides pteronyssinus*, *Blomia tropicalis*, and *Tyrophagus putrescentiae*.

The presence of Blo t 19, a WHO/IUIS allergen, previously identified in *B. tropicalis* could not be confirmed by our genome/transcriptome or GenBank data. Originally, Blo t 19 (GenBank: AHG97583) protein was reported as a putative *B. tropicalis* protein having similarities to the nematode antimicrobial peptide. Our PHMMER search of this peptide did not return any mite matches; all closest matches were nematodes: 81.7% – 85.5% amino acid similarity: *Toxocara canis* KHN74506), *Halicephalobus* sp. KAE9556153, and *Enterobius vermicularis* VDD96638). We therefore consider this previously reported mite allergen as an artifact/contamination.

Group 36 contained two different allergens incorrectly assigned as proteins with the same function (see also Vrtala [13]). The Der p 36 (ATI08932) and Der f 36 (ATI08931) peptides had a C2 domain (1-24 signal peptide, 90-208 C2 domain). The identity of the peptides was 77.6%, while Tyr p 36 had a profilin (alignment region 1-131) structure. Both peptides with C2 domains and profilin are presented in *B. tropicalis*.

In this study we found two Blo t 3 isoforms (Table 1; S5–S6 Figs), but we did not identify in the genome and transcriptome isoform connected to Blo t 3 AAQ24542. However, we identified AAQ24542 protein in proteome [21, 22] (Table 1; S5–S6 Figs). The PHMMER analyses showed that several allergen groups share the same functional domains (Table 1). The allergen Blo t 3 and Blo t 6 both have a trypsin domain. The predicted 3D structures confirmed trypsin-like core structure in JGLJHAHI _02083, JGLJHAHI _03207 and JGLJHAHI _04704 (Fig 2). Except for trypsin-like core structure the rest of the sequences classify them into different groups. The Blo t 3 JGLJHAHI_02083 exhibits similarity to IX factor of blood clotting cascade. Moreover, this protein has been identified only in the bodies, not excrement. The Blo t3 JGLJHAHI_03207 was identified in the excrement and is highly like trypsinogen. The protein structure indicated that trypsinogen is activated by removing of the protein tail (Fig 2). The final one, Blo t 6 JGLJHAHI_04704 exhibits similarity to kallikrein-like serine protease. Both Blo t 5 and Blo t 21 shared the Blo t 5 domain as is confirmed by 2D protein model (Fig 3). The differences is caused by protein tail n Blo t 21.

**Fig 2.**
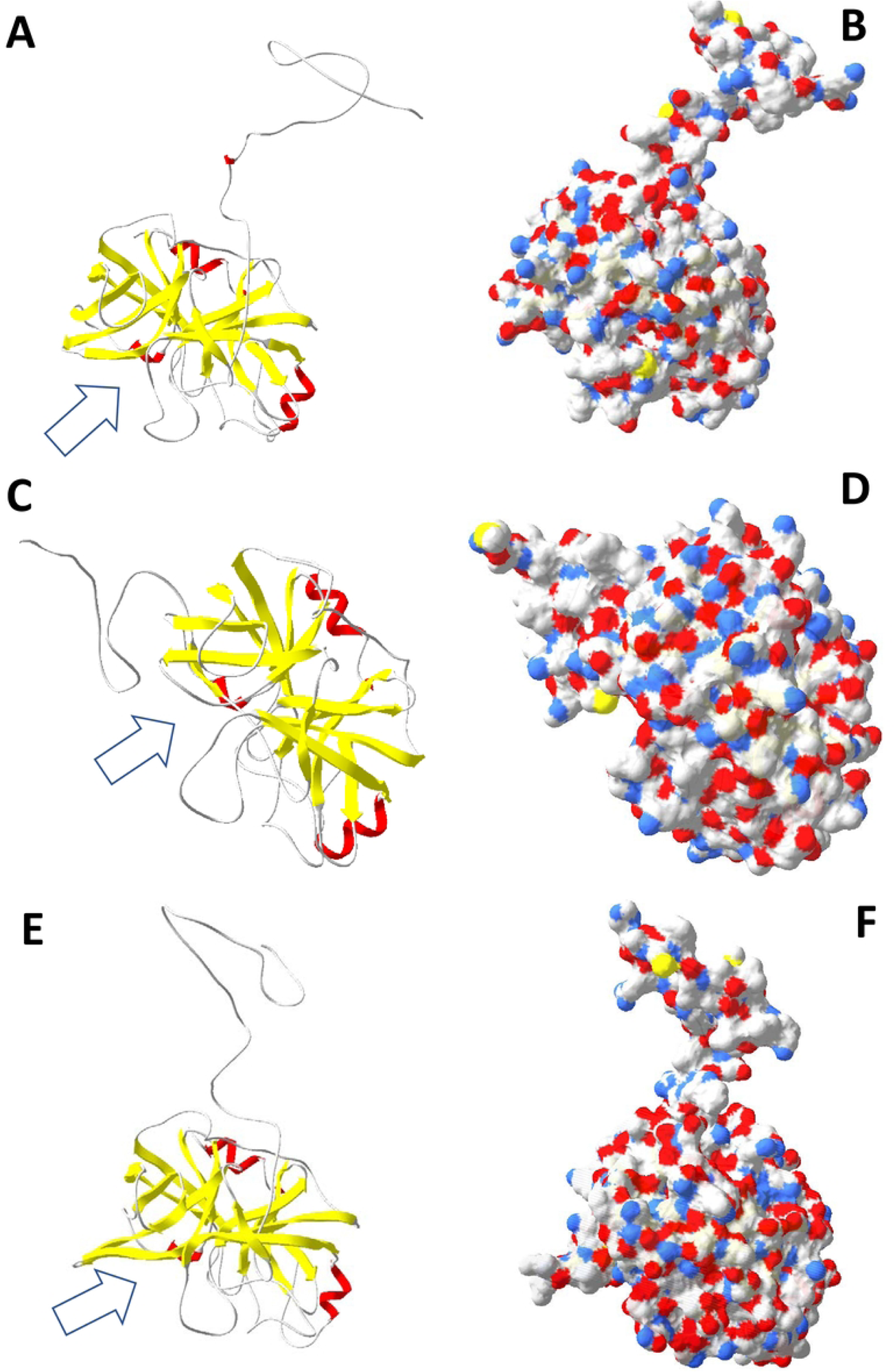
3D models of identified serine proteases IX factor of blood clotting cascade (JGLJHAHI _02083) - **A,B**; trypsinogen (JGLJHAHI _03207) - **C,D**; kallikrein-like serine protease (JGLJHAHI _04704) - **E,F**. The **A,C**, and **D** figures show the backbones. **B,D** and **F** the surface. The positively charged surface is denoted in blue, negatively in red. The active site is indicated by arrow.

**Fig 3.**
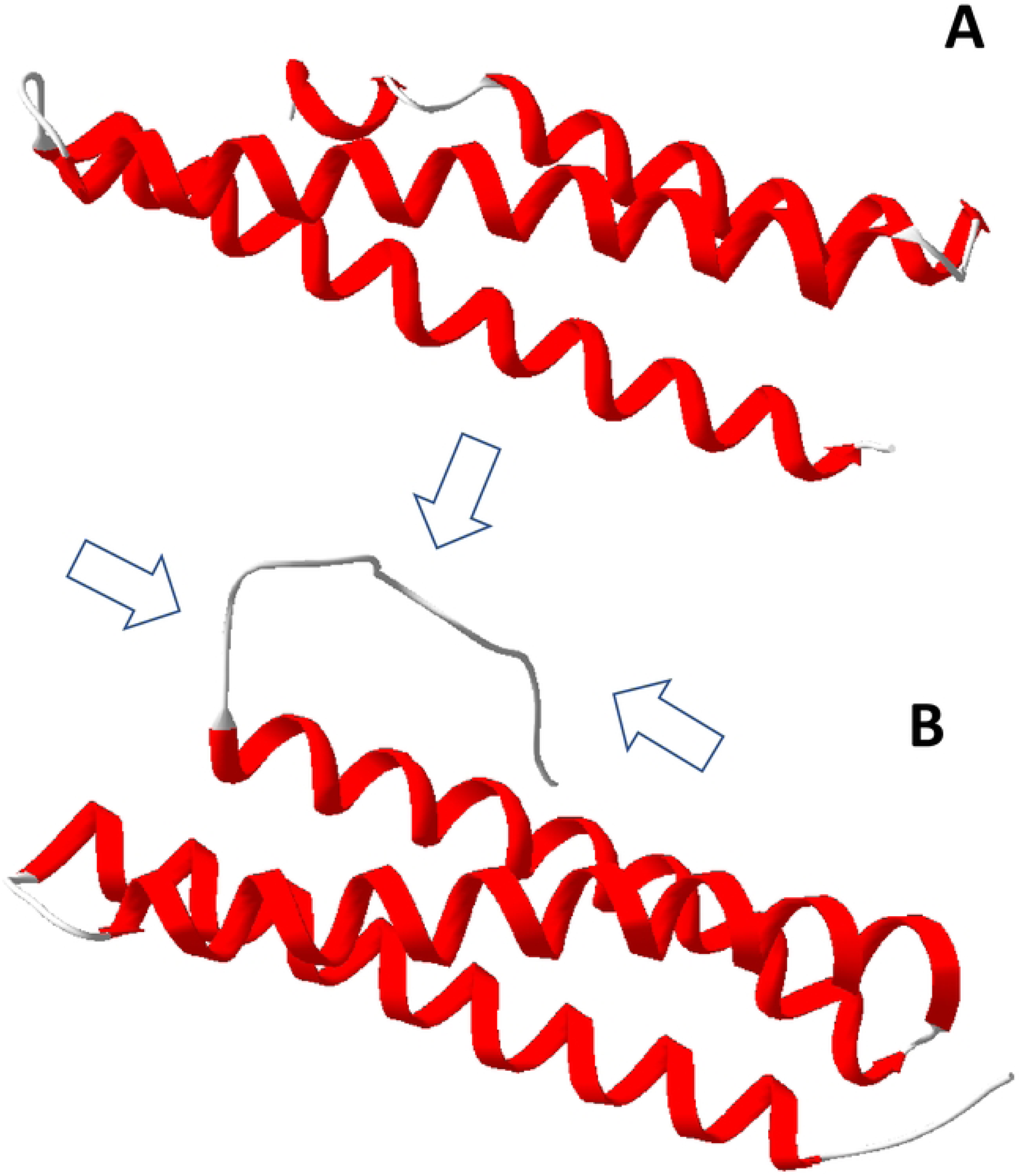
Predicted 2D model of allergen Blot 5 (A) and Blot 21(B). Arrows point to the protein tail.

### Expression of allergen-encoding genes and protein abundance

Our expression analysis identified six highly expressed allergen-encoding genes, Blo t 1, 5, 14, 15, 18, 21 (Fig 4). However, a proteomic analysis identified only some of them as most abundant, Blo t 14, 15, 18 (Fig 5). Proteomes of mite bodies and feces were significantly different in both protein presence/absence (ANOSIM Jaccard, R=0.509, P=0.003) and abundance (ANOSIM Bray–Curtis, R=0.677, P=0.002). The differences (SIMPER analysis) were due to low abundance/absence of several proteins in mite feces: 60S acidic ribosomal protein P2, and Blo t 7, 8, 33, 34 which were represented in mite bodies. In our proteomic samples, alpha-amylase protein (Blo t 4) was highly abundant in both mite bodies and feces, indicating that this enzyme was present in the mite gut and then it was concentrated in feces. Blo t 20 (JGLJHAHI_13643) was absent in fecal pellets but had intermediate abundance in water extracts from feces. Blo t 2 (JGLJHAHI_04625), Blo t 18 (JGLJHAHI_08029) and Blo t 29 (JGLJHAHI_136660) were more abundant in mite bodies than in feces.

**Fig 4.**
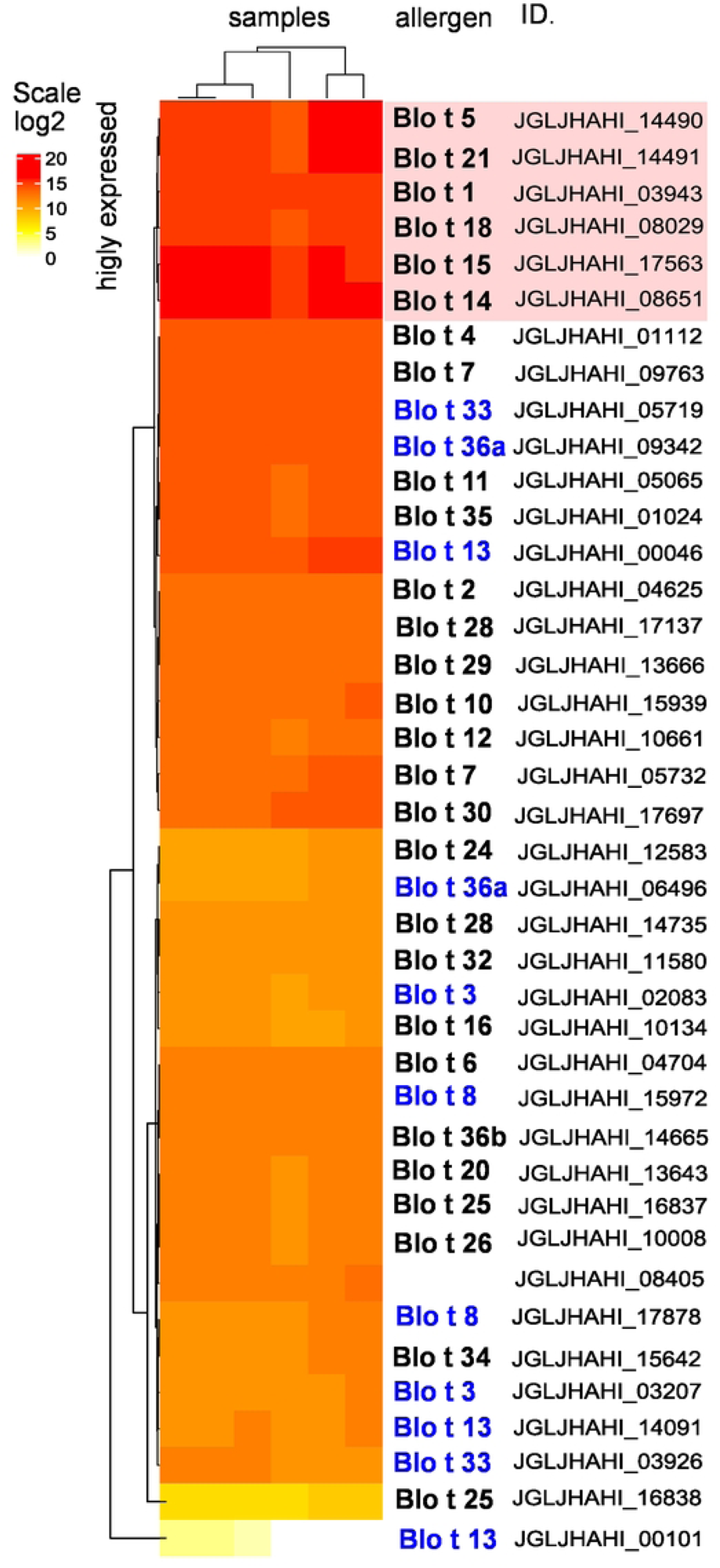
Expression of known and predicted allergen-encoding genes of *B. tropicalis*. Gene expression values were log2 transformed and visualized in a heatmap. The red box indicates allergens of highest expression. In the allergen group names, blue indicates allergens identified or predicted allergen isoforms.

**Fig 5.**
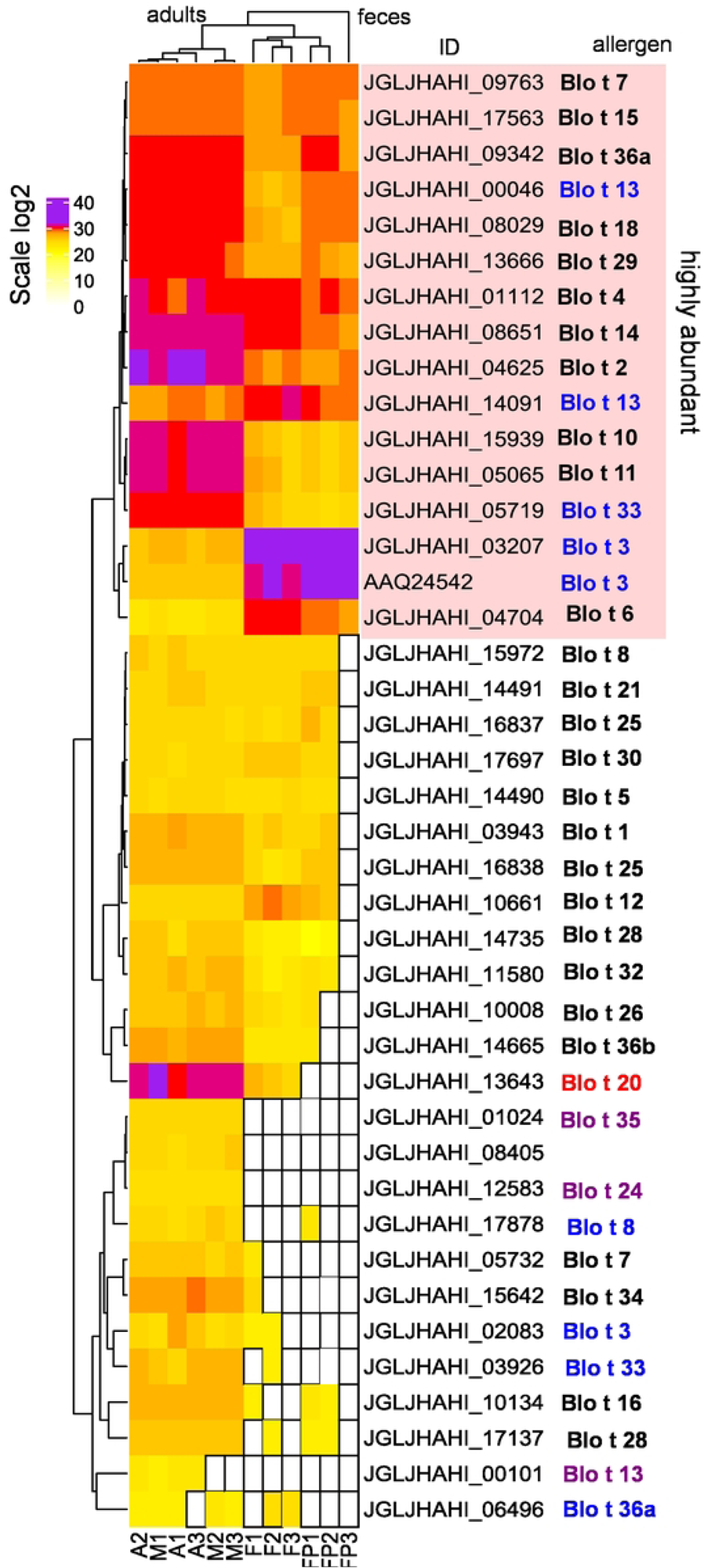
Abundance of allergenic proteins in bodies and feces of *B. tropicalis* (A = adults, M = mixed stages, F = feces, FP = fecal pellets not dissolved in water). Data were log2 transformed. In the allergen group names, blue indicates that an allergen was identified on more isoforms; red indicates that an allergen was not present in fecal pellets (FP), and purple indicates allergens not present in feces (F).

In *B. tropicalis*, group 3 allergen (trypsin) has 3 isoforms (S5 and S6 Figs), but only two of these isoforms were detected in feces at higher abundances as compared to whole mites (e.g., trypsinogen: JGLKHAHI_03207 and AAQ24542). Another isoform, Blo t 3 (IX factor of blood clotting cascade: JGLJHAMI_02083), was nearly absent in feces. Blo t 6 (kallikrein-like serine protease) has a single isoform, which is more abundant in feces than in mite whole bodies. Blo t 13 (lipocalin) has 3 isoforms (S19 Fig). One isoform (JGLKHAHI_14091) predominated in feces, while isoform JGLKHAHI_00046 had almost the same abundance in whole mite bodies and feces, isoform JGLKHAHI_00101 was detected at low abundance in whole mite samples, but not in the feces.

## Discussion

Our multiomic analyses confirmed 13 known groups of *B. tropicalis* allergens (Table 1) and suggested that one previously identified WHO/IUIS allergen (Blo t 19) belongs to a nematode but not to *B. tropicalis* or any other mite. This latter allergen may be a result of contamination by a nematode inhabiting the mite culture medium [23]. Based on structural homology and sequence identity of known WHO/IUIS allergens from pyroglyphid house dust mites *Dermatophagoides* sp. and the mold mite, *T. putrescentiae,* we predicted 16 putative allergens in *B. tropicalis*, groups 14–16, 18, 20, 24–26, 28–30, 32–35, 36 (Table 1). A total of 26 known/predicted *B. tropicalis* allergens, were detected to be common with *Dermatophagoides* and *Tyrophagus* mites. Interestingly, a similar number of 27 candidate allergens was identified previously using a crossed radioimmunoelectrophoresis [11]. Many of these allergens/putative allergens 14 were found in recently published genomic/transcriptomic[22] and proteomic analyses [20].

Our study showed that Blo t 3, 8, 13, 33 36a (peptides with C2 domain) occurs in more isoforms, however we did not confirm isoforms for Blo t 2. Blo t 2 allergen protein was abundant in mite bodies as well as fecal pellets (Fig 4). We found only a single isoform, but contrasts with Reginald at al. study [24] which reported nine Blo t 2 isoforms having 1–3 amino acid substitutions [24]. As these isoforms were obtained via RT-PCR/cloning and were not verified using direct DNA sequencing [24], it is unknown whether these substitutions represent the natural variation of the protein, polymerase errors, or a mix of the two. The previous proteomic study revealed 3 isoforms of different protein profiles in *B. tropicalis* extracts.[20] Lipocalin Blo t 13 is minor allergen [12, 25] having 3 isoforms (our data; Some of these isoforms had higher abundances in mite feces (Fig 4) as noted previously for *T. putrescentiae* [18].

Blo t 14 allergen (vitellogenin) [20] showed higher levels in both whole mite bodies and feces. In *D. pteronyssinus* the allergen has been described as an apolipophorin-like group 14 allergen, with predicted hydrophobicity and lipid-binding activity, and inducing high levels of IgE responses and T-cell stimulation [26]. This is an exoskeleton-bound protein having much higher abundance in females of than in males *D. farinae* [23]. Contradiction with “exoskeleton-bound”, vitellogenins are yolk protein precursors that are produced at extra-ovariole sites and taken up by the growing oocytes [27]; the presence of vitellogenin in feces is thus surprising and protein structure needs further attention. However it is suggested that the can bind several microbial ligands as LPS, lipoteichoic acid or β-glucan due to the vitellogenin domain (as reviewed Jacquet [28]).

Putative Blo t 20 allergen protein (arginine kinase[29] or ATP:guanido phosphotransferase (our PHMER analysis)) is relatively abundant in mite bodies but has low or zero abundance in feces. Der p 20 has 40% IgE reactivity in non-hospitalized patients [30], and Der f 20 showed 7% IgE reactivity to patient sera [31, 32], indicating that in pyroglyphid mites this allergen has a moderate to low medical importance. However, arginine kinase is a major allergen in crustaceans that may cause asthma and fatal anaphylaxis in fishermen and processing plant workers [33]. Allergenic properties of putative Blo t 20 are unknown.

Protease allergens (Blo t 3 trypsin and Blo t 6 chymotrypsin) were predominant proteins in mite feces. Similarly, group 3 and 6 proteins were highly abundant in the mold mite *T. putrescentiae* feces [17–19] but in *D. pteronyssinus*, only chymotrypsin ^17,18^. Previous observations suggest that in culture extracts of *B. tropicalis*, no trypsin or chymotrypsin enzymatic activity could be detected [34] or these enzymes have been found at low abundance [20]. However, another study based on design using inhibitor and specific substrate for enzyme identified trypsin, elastase, chymotrypsin, kallikrein, C3/C5 convertase, and mast cell protease [35]. Our results showed that two isoforms of Blo t 3 differ by their protein structure and body localization; i.e. body localized Blot 3 JGLJHAHI_02083 of similarity to IX factor of blood clotting cascade; and feces associated trypsinogen JGLJHAHI_03207. The last Blo t 3 (AAQ24542) isoform was obtained by amplification of isolate RNA and confirmed by DNA sequencing [36]. The absence the transcriptome and presence of *B. tropicalis* proteome in this and previous study [20] indicated posttranslational modification. Surprisingly Blo t 6 JGLJHAHI_04704 exhibits similarity to kallikrein-like serine protease confirmed presence of kallikrein enzymatic activity of *B. tropicalis* extract [35]. These results can be reconciled as enzyme inactivation may occur in the mite bodies [37] or the expression of these enzymes were affected due to the presence of intracellular bacterial symbionts, i.e. *Cardinium* and *Wolbachia* [38].

Blo t 5 and 21 have been reported as the major allergens of *B. tropicalis* [10, 13]. Our analyses showed that both proteins contained a signal peptide and share a Blo t 5 domain. The majority (>75%) of sensitized patients were co-sensitized to both Blo t 5 and Blo t 21, which is expected based on their structural similarity[39] (S51 Fig). Both of these allergens were found in the midgut and inside fecal pellets [39], as independently suggested by our gene expression and to our proteomic analyses.

In conclusion, we show that the medically important mite *B. tropicalis* has a large set of putative allergens (n=15) predicted by our omics analyses (genome, transcriptome, proteome) and the use of carefully verified comparative data on known allergens. Although, the clinical importance of these putative allergens is be verified in the future, a similar bioinformatics approach was successful for allergen discovery in the mold *T. putrescentiae* [40]. It is, therefore, expected that a large portion of our predicted putative allergens will be of medical significance. Our study also provides accurate quantification of immunogenic proteins in both mite whole bodies and fecal pellets, the two important mite-related components having distinct allergenic profiles. For this task we used a shotgun proteomics approach with an advanced Tribrid Orbitrap Fusion spectrometer and contrasted our results with gene expression data. We show that expression levels of allergen-encoding genes may not be strictly correlated with the actual allergenic protein abundance. Effective strategies for identification and prediction of new allergens include high-throughput sequencing, proteomics, transcriptomics and genomics; however, most accurate results are achieved when these technologies are integrated in а so called *omics* approach [23, 38, 41–45]. Moreover, label-free or other quantitative approaches enable the estimation of mite allergen abundance in proteome profiles [23, 38].

## Material and methods

### Sample of mites and feces

Laboratory culture of *B. tropicalis* was maintained as described previously [46] at the Crop Research Institute, Prague, Czechia. Mites were mass reared in tissue culture flasks on a house dust mite diet (HDMd) in a large population (approximately 4 weeks) when the diet is almost consumed. Detailed rearing and harvesting methodology is described elsewhere [47]. The adult or juvenile mites were collected from the plugs and surfaces of the flasks with a brush and placed in sterile microfuge tubes. “Pure” feces were obtained from the flasks after the mites and remnants of their bodies or eggs were removed and collected [18].

The fresh weights of the mites or their feces were in the range of 30–40 mg using a microbalance (Mettler-Toledo). For transcriptomics, mites were surface cleaned by placing them in 100% ethanol, followed by vortexing for 5 s and centrifugation at 13,000 × g for 1 min. The supernatant was replaced with a 1:10 bleach solution containing 5% sodium hypochlorite, and the samples were then mixed by vortexing for 5 s and centrifuged at 13,000 × g for 2 min. The last step was washing twice with dd H_2_O to remove residual bleach. Surface sterilization was performed on ice. The rearing diet had the same weight as the feces. The rearing diet and feces were not cleaned and were stored directly in a ultra-cold freezer (−80° C). We collected five biological replicates of mites for RNA sequencing and 1 sample of mites for genomic sequencing. Every sample contains approximately 1,000 mites.

For proteomic analyses, four sample types, each with three biological replicates, were prepared: (i) 1000 individually collected adult mites; (ii) pools of mites of different ages, including eggs; (iii) water extracts of feces; and (iv) detergent-buffer extracts of the remaining pellet of the water extract.

### Sample processing

Homogenization of mites was performed on ice. All samples were homogenized for 30 s in a glass tissue grinder (Kavalier glass, Prague, Czechia) in 500 μl of lysis buffer from the kits. DNA was extracted from homogenates after overnight incubation with 20 μl of proteinase K at 56°C using the QIAamp DNA Micro Kit (Qiagen, Hilden, Germany, cat. No. 56304) following the manufacturer’s protocol for tissue samples. The concentration of the extracted DNA samples was quantified using the Qubit^®^ dsDNA HS Assay Kit (Life Technologies), and the quality of the DNA was determined using a NanoDrop 2000 spectrophotometer (Thermo Fisher Scientific). The average size of gDNA was determined using E-Gel SizeSelect 2% agarose gel (Invitrogen) with a 1 kb ladder. The samples were sheared using a Covaris G-tube (Covaris Inc.). The average size of the sheared DNA was determined using an Agilent 2100 Bioanalyzer (Agilent Technologies).

RNA isolation was performed using the NucleoSpin RNA kit (catalog no. 740984.50; Macherey-Nagel, Duren, Germany) according to the manufacturer’s instructions, with the following modifications: homogenized samples were centrifuged at 2,000 × g for 3 s, and DNA was degraded by DNase I at 37°C according to the manufacturer’s protocol (Riboclear plus, catalog no. 313-50; GeneAll, Lisbon, Portugal). RNA quality was evaluated using a NanoDrop (NanoDrop One; Thermo Scientific, Waltham, MA, USA) and an Agilent 2100 Bioanalyzer (Agilent Technologies, Santa Clara, CA, USA), and samples altogether with DNA sample were shipped on dry ice to the MrDNA laboratory (Shallowater, TX, USA) for downstream processing and sequencing.

For protein analyses, the mite body samples (i and ii) were further homogenized in a Potter-Elvehjem homogenizers with a Teflon pestle after the addition of 100mM (1 mL/1000 mites) triethylammonium bicarbonate buffer (TEAB; Cat No. 17902, Fluka, Sigma‒Aldrich, St. Louis, MO, USA) containing 2% w/v sodium deoxycholate (SDC; Cat No. 30970, BioXtra, Sigma‒Aldrich). Water fractions of feces (iii) were obtained in the tissue cell culture flasks as previously described [17–19]. Briefly, 10 mL of ice-cold (4°C) Nanopure water (Thermo, Waltham, MA, USA) was added to the feces-coated flasks. The flasks were placed in a box with ice and mixed on a ProBlot Rocker (Labnet International, Woodbridge, NJ, USA) for 10 min laying on the side at 100 RPM, and the extract was transferred to 50 mL sterile flasks. The second side of the feces-coated flask was extracted with 10 mL of sterile water, and the extract was then added to the 50 mL flask. The third extraction in the flask was performed with 10 mL of water, which was thoroughly mixed in the flask. In total, 30 mL of the extract obtained from a flask was centrifuged at 10,000xg for 15 min at 4°C (Thermo). The supernatant was divided into 3 equal portions, and portions of 1 mL were collected for total protein quantification. The contents were lyophilized in a PowerDry LL3000 lyophilizer (Thermo Fisher Scientific). The pellets of feces (iv) used for proteomic analysis was obtained by processing the pellet after water extraction. 3 mL of 100 mM TEAB with 2% SDC was added to the pellet that remained from approximately 20 mL of the supernatant, and the content in the centrifuge tube was thoroughly mixed by vortexing plus 10 min of mixing on the rocker on ice. The procedure was repeated 3 times. The resulting sample was centrifuged at 10,000xg for 15 min at 4°C (Thermo Fisher Scientific), and the supernatant was collected for further analysis. All the samples were stored at −80°C until use.

### Sequencing and assemblage

For Illumina DNA sequencing, the libraries were prepared using a Nextera DNA Flex library preparation kit (Illumina) following the manufacturer’s user guide. Fifty nanograms of DNA was used to prepare the libraries. The samples simultaneously underwent fragmentation and addition of adapter sequences. These adapters were utilized during limited-cycle PCR in which unique indices were added to the samples. The libraries were then pooled in equimolar ratios of 0.6 nM and subjected to paired-end sequencing for 500 cycles using a NovaSeq 6000 system (Illumina).

For PacBio sequencing, 250 ng of the sheared DNA was employed as the input for library preparation using the SMRTbell Express Template Prep Kit 2.0 (Pacific Biosciences). During library preparation, each sample underwent DNA damage and end repair as well as barcode adapter ligation. To reduce the number of smaller DNA fragments (< 3 kb), an additional bead purification step was performed on the SMRTbell libraries using 40% diluted AMPure PB Beads. Each library was then sequenced using a 10-hour movie time on a PacBio Sequel system (Pacific Biosciences). The SMRT Link Circular Consensus Sequencing workflow (SMRT Link v.9.0.0, CCS) was used to combine multiple subreads from the same molecule to generate a highly accurate consensus sequence. The sample metadata and the whole genome sequence were deposited in GenBank as part of project PRJNA625856.

Concentration of total RNA was determined using the Qubit^®^ RNA Assay Kit (Life Technologies). Poly-A selection and library preparation were performed by using KAPA mRNA HyperPrep Kits (Roche) following the manufacturer’s instructions. A total of 1000 ng RNA was used to prepare the library. Following library preparation, the final concentration of the library was measured using the Qubit^®^ dsDNA HS Assay Kit (Life Technologies), and the average library size was determined using an Agilent 2100 Bioanalyzer (Agilent Technologies). The library was then diluted to 0.6 nM and subjected to paired-end sequencing for 500 cycles using the NovaSeq 6000 system (Illumina). The samples are deposited in GenBank as project PRJNA599071.

Illumina reads were trimmed in Trim Galore [48], corrected in fastQC [49], aligned with the PacBio reads in hybrid SPADES v 3.14 [50, 51] for DNA-based reads and rnaSPADES for RNA-based reads [50].Transcriptome sequences were annotated by Prokka [52], and predicted proteins were identified by a KEGG analysis in GhostKoala [53]. Reads were mapped onto assemblage genome or transcriptome using Bowtie2 [54, 55] and Minimap2 [56] for long sequences. Then the assemblage genomes were improved by Pilon [57]. The GFF file from genome and transcriptome was prepared by EXONERATE [58]. MAKER [59] and FUNANNOTATE [60] on the Galaxy server [61]. The expression was done in CLC Worbench 22 (Qiagen, Venlo, Netherlands) using total number of mapped reads as expression value.

### Protein analyses

Samples for label-free mass spectrometry were processed and further analyzed using a nanoLC‒MS/MS system employing an Orbitrap Fusion Tribrid mass spectrometer (Thermo Fisher Scientific) as previously described [23, 38]. Mass spectrometry data were evaluated in MaxQuant version 2.2.0.0 using label-free quantification (LFQ) algorithms [62, 63] and the Andromeda search engine [64]. The key criteria used in data analyses were a false discovery rate (FDR) of 0.01 for proteins and peptides, a minimum length of 7 amino acids, carbamidomethyl as a fixed modification, and variable modifications of N-terminal protein acetylation and methionine oxidation. This data were searched using a custom database with our transcriptome assemblies of *B. tropicalis*. In addition, we included a GenBank genomic sequence of *B. tropicalis* (NCBI: 16792seq; 06 Feb 2023), and sequences of possible associated contaminants and microorganisms: *Aspergillus* (UniProtKB: 5439 seq 07 Feb 2023), *Staphylococcus kloosii* (UniProtKB 4243seq: 07 Feb 2023), *Staphylococcus rev* (UniProtKB: 13547seq, 07 Feb 2023), *Staphylococcus xylosus* (UniProtKB: 12883, 07 Feb 2023), and yeast (UniProtKB: 21667seq, 07 Feb 2023). The data were processed in Perseus version 2.0.7.0 [65].

### Allergen identification

Sequences of allergens were downloaded from the WHO/IUIS database [4] and subjected to p BLAST searches against the GenBank database using BLASTp [66]. The closest search results for *B. tropicalis* and other mites were added to extend mite WHO/IUIS allergen database. Then, the predicted proteins from *B. tropicalis* were screened against the allergen database using HMMER [67] on the Galaxy server. Predicted sequences with the highest scores were selected for analyses and compared to the proteome identification (see above).

In the next step we performed structural alignment on the TCofee server [68, 69] for each allergen group. For homologous sequence clusters, phylogenetic trees were inferred from in PhyML [70]. The trees were visualized and edited in FigTree v.1.4.4 [71], alignments were visualized in ESPrint [72]. Sequence identities were calculated in EMBOSS Water using the Smith–Waterman algorithm [73]. The protein structure, including the signal peptide, was predicted based on comparison to reference proteins in the EMBL database [74].

Trypsin identification was based on BLASTp we searched for the homologous pdb structures. Then JGLJHAHI _02083, JGLJHAHI _03207 and JGLJHAHI _04704 were aligned to the structures with the results of BLASTp searches using CLUSTALlX 2.0 [75]. The protein alignment was then used as an input for MODELLER 10.2 software [76, 77] for 3D structure prediction. The validity of the resulting models was checked using PROCHECK software(G-score) [78], and the model with the best valid score was chosen for further investigation. This model was optimized by running 10 steepest descent energy minimization steps followed by 10 conjugated gradient, 20 steepest descent, 10 conjugated gradient and finally 10 steepest descent steps (GROMOS 96 force-field) [79]. The validity of the final model was again checked by PROCHECK.

### Data analyses

Expression and protein heatmaps were created in R-4.2.2 software [80] using the Complex-Heatmap package [81, 82]. Protein profiles of feces and mite bodies were compared using analysis of similarities (ANOSIM). Two distance matrices were chosen: the Jaccard index (presence/absence of proteins), and the Bray–Curtis index (protein hit intensity). To identify allergens/predicted allergens responsible for the differences between the profiles, we used similarity percentages (SIMPER). The calculations for all of these analyses were performed in PAST 4 [83].

## Acknowledgments

The authors thank Marta Nesvorna, Julie Chalupnikova, and Martin Markovic (all Crop Research Institute, Prague, Czechia) for technical help. JH, BS, TE were supported by project LUAUS23082 of the Czech Ministry of Education, Youth and Sports. PBK was supported by the Ministry of Science and Higher Education of the Russian Federation within the framework of the Federal Scientific and Technical Program for the Development of Genetic Technologies for 2019-2027 (agreement №075-15-2021-1345, unique identifier RF 193021X0012).

The authors acknowledge the support of the Freiburg Galaxy Team: Person X and Bjorn Gruning, Bioinformatics, University of Freiburg (Germany) funded by the Collaborative Research Centre 992 Medical Epigenetics (DFG grant SFB 992/1 2012) and the German Federal Ministry of Education and Research BMBF grant 031 A538A de.NBI-RBC.

## Author contributions

JH, ET, PBK, QH BS and SV – scientific writing, JH – detailed data evaluation and preparation of figures and tables, KH, PT, and ET – proteomic analysis performance, SD – genome and transcriptome sequencing, BS – protein analyses, ET – experimental design and all authors commented the final version of draft.

## Data availability

Sequences and samples obtained during this work have been submitted to the NCBI Respiratory (SUB7787303). The accession number for the raw nLC-MS/MS runs reported in this paper is MassIVE MSV000091854 (doi:10.25345/C5VD6PF6P) or PXD041972. Furthermore, we provide the entire “combined/txt” folder from MaxQuant data processing, and the protein databases used for the search for download.

